# Efficient *De Novo* Assembly and Recovery of Microbial Genomes from Complex Metagenomes Using a Reduced Set of *k*-mers

**DOI:** 10.1101/2024.06.08.598064

**Authors:** Hajra Qayyum, Amjad Ali, Masood Ur Rehman Kayani

**Author notes:** To whom correspondence should be addressed: Amjad Ali, and Masood Ur Rehman Kayani.

## Abstract

In recent years, the analysis of metagenomic data to recover unculturable microbes has revolutionized microbial genomics by rapidly expanding the reference genome catalog. Central to this, are the computational approaches of *de novo* assembly and genome binning that enable large-scale reference-independent recovery of microbial genomes from the metagenomic sequencing data. Despite the advancements in bioinformatics approaches to address the computational challenges inherent to these tasks, the limitation of computational resources continues to be a significant barrier to harvesting the full potential of these techniques. Consequently, there is a stressed need to devise strategies involving the fine-tuning of the employed parameters for the effective utilization of the available metagenomic tools. As most of the available metagenome assembly tools are based on the *de Bruijn* graph framework that relies on a parameter *k*, selecting an appropriate subset of *k*-mers has become a common approach in bioinformatics for efficient computations. In this study, we propose a reduced set of *k*-mers, optimized to strike a balance between computational efficiency and the quality of the high- and low-complexity metagenome assemblies. Utilizing this set of *k*-mers with MEGAHIT reduces the metagenome assembly time by half compared to the default set, thus greatly reducing the associated computational cost. In addition, it also brings the promise to improve large-scale genome binning studies that adopt this set in the future as we observed an increase in the total number of the recovered genomes as well as obtained higher proportions of high- and medium-quality genomes recovered from the reduced *k*-mers-based metagenome assemblies.

## INTRODUCTION

With the advent of next-generation sequencing (NGS), there has been a rapid surge in metagenomic studies, substantially advancing our knowledge of unculturable microbial majority and their association with health and disease. Recently, genome-centric metagenomic analyses have contributed to the recovery of thousands of high-quality microbial genomes from the human microbiome [1], [2] most of which serve as the first genomic references to previously uncharacterized microbial species. These accomplishments underscore the transformative capabilities of NGS. Central to such analysis is the intricate task of assembling millions of short reads (typically 100-150 bp in length) generated by common NGS platforms into larger contiguous segments (termed contigs) through a computational process called metagenome assembly [3]. These contigs are then clustered into metagenome-assembled genomes (MAGs) by yet another computational process known as genome binning, followed by downstream analyses based on the objectives of the individual studies.

The metagenome assembly process is typically *de novo* since most of the microbes do not have a representative in the genome databases. Dedicated metagenome assemblers have been developed for this purpose, which include the MEGAHIT [4], metaSPAdes [5], IDBA-UD [6], and many others [7],[8],[9]. These tools employ the Overlap-Layout-Consensus or *de Bruijn* graph (DBG)(*k*-mers-based) approach, with the latter being the most prevailing and widely adopted algorithm [10]. Regardless of the strategy and the employed tools, the metagenome assembly faces multiple challenges that impede its efficiency as well as accuracy. Computational efficiency becomes a major bottleneck in metagenome assembly, especially for larger and more complex metagenomes such as the human gut microbiome [11], [12]. For instance, Pasolli et al. (2019) reported that several human gut microbiome samples required >1TB of computer RAM for their assembly using metaSPAdes [1]. This makes metagenome assembly of such samples practically impossible, especially in settings with limited computational resources.

While there are several computationally optimized frameworks such as Matchtigs [13], MegaGTA [14], and SparseAssembler [10] designed for memory-efficient metagenomic assembly, there are also time-optimized solutions like ScalaDBG (patch to IDBA-UD), designed to accelerate the metagenome assembly process by exploiting the parallel DBG construction for multiple *k*-mers [15]. However, it is important to note that these approaches either suggest a framework or optimization to the existing DBG algorithm and do not address the optimization of the employed parameters in existing metagenomic tools. These computationally less-intensive approaches can be deployed in such cases, but it should be acknowledged that they are not devoid of complexities and challenges.

As mentioned above, most of the current metagenomic assemblers are DBG-based [7], [16], where the nodes of the graphs represent sub-strings of the query sequences of length *k* (*k-*mers) interconnected by edges. Hence, the selection of appropriate *k*-mers becomes an integral aspect of obtaining a high-quality metagenome assembly [10]. Practically, the choice of the *k*-mers is based on previous experiences with similar datasets or exploiting brute force by experimenting with different values of *k*-mers; however, these strategies can take a substantially longer time for complex metagenomes, even with the fastest available metagenome assemblers [17]. Some studies have also adopted the strategy of informed choice of *k*-mers by building abundance histograms for possible values of *k* and then comparing the results; however, this approach is again limited by time, as constructing a histogram just for one *k* is time-consuming [18]. Given the constraint of limited computational resources and metagenome complexity, there is an immediate need to design a fine-tuned *k*-mers set to maximize the utility of existing bioinformatics tools for speeding up computations.

In this study, we tested various *k*-mers sets for the assembly of human metagenomes as well as for MAG recovery, to provide an optimal set of *k*-mers capable of optimally performing for metagenomes of variable complexities. The proposed set of *k*-mers achieved assembly in significantly reduced computational time and without trading off the quality of the assemblage and recovered MAGs. We believe that this computationally efficient solution will empower the scientific community, especially those with limited computational resources, to effortlessly conduct high-quality metagenomic analyses in minimal processing time.

## METHODOLOGY

### Metagenome Samples and Preprocessing

Here, we used publicly available metagenomes from the Human Microbiome Project Phase III [19]. Specifically, we recruited 70 gut and 30 anterior nares samples (further referred to as skin samples) as representatives of high and low-complexity metagenomes, respectively. The accession numbers and brief metadata of these samples are provided in Supplementary Table I. These metagenomes were retrieved from the NCBI SRA database using the prefetch program (v3.0.2) provided in NCBI’s SRA Toolkit (https://github.com/ncbi/sra-tools). Following retrieval, the fasterq-dump (v3.0.2) was used for data conversion in the FASTQ format. Raw FASTQ files were then subjected to quality assessment and preprocessing using a combination of FastQC (v0.12.1) (https://github.com/s-andrews/FastQC) and fastp (v0.23.2) [20]. Using fastp, low-quality bases (Q<20), adaptor sequences, and duplicate reads were removed (using parameters: *-q, -- detect_adapter_for_pe, and --dedup,* respectively) from the raw reads to obtain high-quality reads, which were used in the downstream analyses.

### De Novo Metagenome Assembly and Quality Assessment

Preprocessed reads were *de novo* assembled using a *de Bruijn* graph-based assembler, i.e., MEGAHIT (v1.2.9) [4]. For this, we created two sets of *k*-mers, further referred to as the “reduced” and “extended” sets, since they are based on the reduction or extension of the *k*-mers used in default settings of MEGAHIT. The reduced set included *k* values of 31, 59, 87, and 115 (MEGAHIT option: *--k-list 31, 59, 87, 115*). In contrast, the extended set included *k* values ranging from 21 *to* 119 with a step size 6 (*--k-min 21*, *--k-max 119,* and *--k-step* 6).

Next, we *de novo* assembled each metagenome using the three sets of *k*-mers, i.e., default, reduced, and extended. All parameters were used with their default settings except the minimum contig length, which was increased to 1500 bp (--*min-contig-len* 1500) in all three sets. For a preliminary assessment of the metagenome assemblies, we computed the general statistics, including the total number of contigs, total assembly length, length of the largest contig, average contig length, and N50 length using QUAST (v5.2.0) [21]. To further evaluate the quality of the assembled metagenomes, we used DeepMAsED (v0.3.1), a deep learning-based approach that can identify misassembled contigs in the absence of reference genomes [22], with its default settings. For this, we selected 9 metagenomes that were best assembled using the reduced *k*-mers set. Collectively, the 27 assemblies for these metagenomes were contained in ∼300,000 contigs (Supplementary Table I). Misassemblies were calculated by DeepMAsED in two steps: first, feature tables were generated for all the assemblies (*DeepMAsED features*), and in the second step feature tables were used for predicting the misassembled contigs (*DeepMAsED predict*).

### Recovery and QC of MAGs Across Three Sets of k-mers

We then sought the recovery of MAGs from each assembly of metagenomes from the three different settings of *k*-mers. To this end, we first mapped the clean reads on respective assemblies using the Burrows-Wheeler Aligner (v0.7.17-r1188) (*bwa mem* command) [23] with default settings. Alignments were then sorted using SAMtools (v1.17) [24] and indexed (using *samtools sort* and *index*, respectively). Subsequently, the *jgi_summarize_bam_contig_depths* function from MetaBAT2 (v.2.12.1) [25] was used to calculate the contig depths in each assembled metagenome. Next, we performed the recovery of MAGs using two of the most commonly used genome binning methodologies, i.e., MetaBAT2 and MaxBin 2.0 (v2.2.7) [26], using their default parameters.

For determining the quality of the recovered MAGs in terms of completeness and contamination levels, we used the lineage workflow (*lineage_wf*) from CheckM (v1.2.2) [27]. For filtering low-quality MAGs, we adopted similar standards as defined by the Minimum Information about Metagenome-Assembled Genome (MIMAG) [28]. In brief, medium-quality (MQ) MAGs were defined as MAGs with completeness ≥50% and contamination <10%. In contrast, a MAG was defined as high-quality (HQ) if its completeness level was >90% and contamination was <5%. All other MAGs were considered low-quality and discarded from further analyses. In addition to the completeness and contamination levels, other metrics for the MQ and HQ MAGs (genome length, number of contigs, largest contig size, and N50 length) were estimated using QUAST (v5.2.0) [21].

### Validating the Utility of the Reduced Set of k-mers

For validating the utility of the reduced set of *k*-mers in efficient metagenome assembly and MAGs recovery, we selected 50 additional gut metagenomes (Supplementary Table II) published by Vincent et al. [29]. These metagenomes are among the massive pool of human metagenomes recruited by Pasolli et al. [1] for the successful recovery of >150,000 MAGs. During their work, these samples were assembled using MEGAHIT (with default parameters), whereas MAGs were recovered using MetaBAT2. We subjected these samples to *de novo* assembly using the reduced set of *k*-mers and recovered MAGs using the default parameters of MetaBAT2. Due to their longer read length (i.e., 150bp), we added another *k* to our reduced *k*-mers list, hence, the reduced *k*-mers set for these samples was: 31, 59, 87, 115, and 143. We further assessed the quality of the recovered MAGs using CheckM, as described in the previous section. To enable comparison, we downloaded the metagenome assemblies, and corresponding MAGs for these samples from http://segatalab.cibio.unitn.it/data/Pasolli_et_al.html (accessed: August 15, 2023). In addition, we retrieved the CheckM reports for the MAGs from the supplementary information of Pasolli et al. [1]. The standard metrics for the assemblies generated using the two datasets were computed using QUAST. For classifying MAGs into HQ and MQ groups, we adopted the criteria defined by Pasolli et al. [1]. Briefly, MAGs in this comparison were considered HQ if their completeness was >90 and contamination was <5, whereas MQ MAGs had a completeness level of ≥50 and contamination <5.

### Statistical Analysis

The time taken by each analysis step (i.e., from assembly to MAG QC) was calculated using the *time* command in Linux. The statistical significance for differences between different categories was calculated by employing the non-parametric Wilcoxon rank sum test [30] in R (v4.3.0). Differences were considered statistically significant for *P* < 0.05.

## RESULTS

### Computationally Efficient Metagenome Assembly with the Reduced Set of k-mers

In this study, we tested three sets of *k*-mers (default, reduced, and extended) to assemble metagenomes of high- and low-complexities using MEGAHIT and recover microbial genomes from the respective assemblies. Our results indicate that the reduced set of *k*-mers outperformed the default and the extended *k*-mers sets by assembling the recruited metagenomes in significantly reduced time. Assembly of the gut samples using the reduced *k*-mers took ∼29±6.97 minutes in contrast with 42.5±15.94 and 84.5±24.62 minutes taken by the default and extended sets, respectively (Fig.1(a)). The pairwise comparative analysis of the computational time taken by the metagenome assemblies for the three categories indicated highly significant differences (*P* <0.0001) using the Wilcoxon rank sum test (Fig. 1(a)).

**Figure 1:**
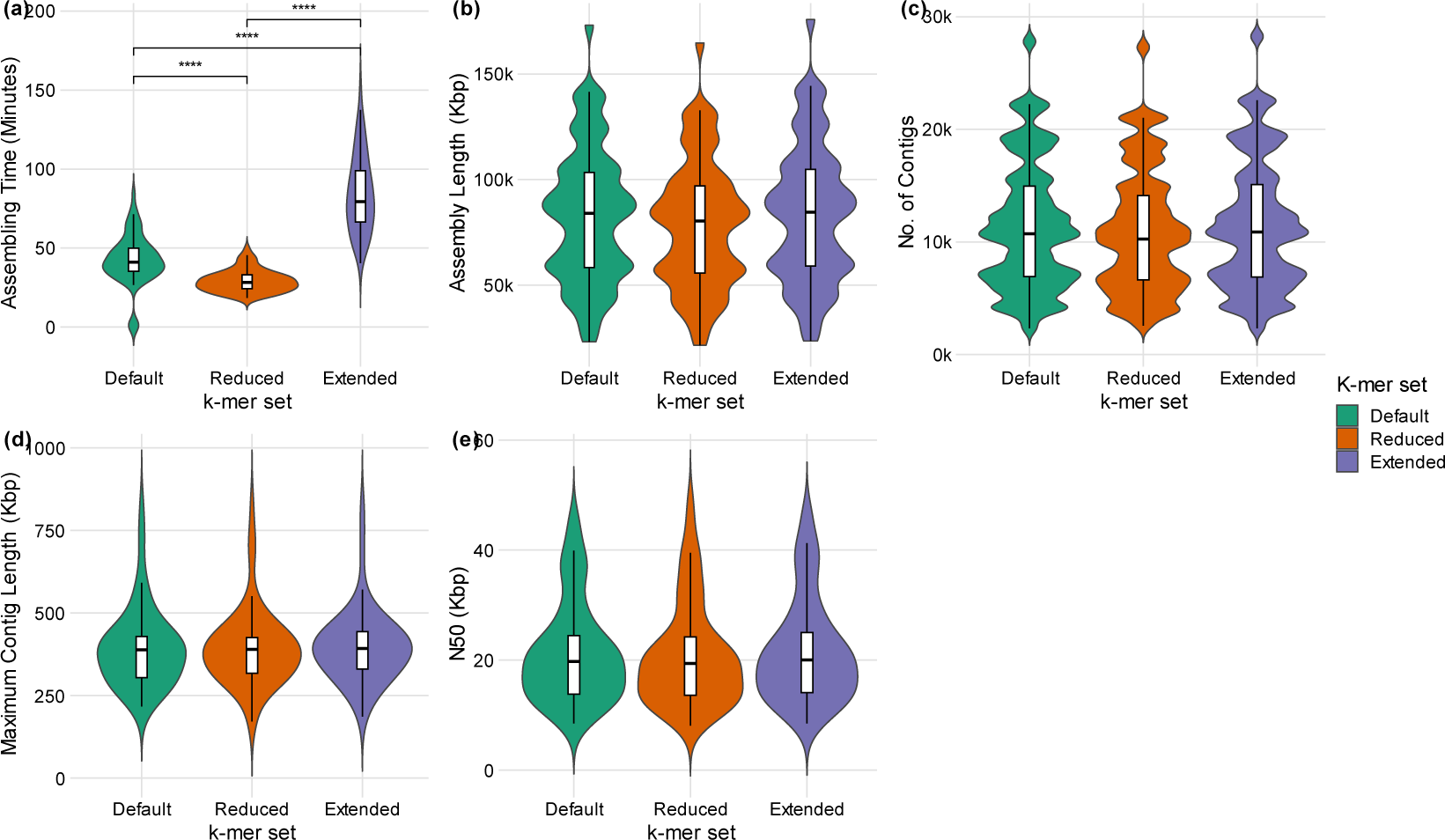
Comparative analysis of the assembly metrics of human gut metagenome samples between default, reduced and extended *k*-mer sets. (**A)** Comparison in the processing time of the assembly performed using different *k*-mer sets. **(B)** Pairwise comparison between the number of contigs generated by the three *k*-mer sets. **(C)** Comparison between the N50 lengths of the assembly attained by different *k*-mer configurations. **(D)** Comparison between the total assembly length achieved by the different *k*-mers sets. **(E)** Comparison between the maximum contig length acquired by the three *k*-mer sets. **Note:** In the figure, asterisks represent the p-value as determined by the Wilcoxon rank sum test (** = <0.01, *** = <0.001 and ****= p-value <0.0001)

To assess the impact of our reduced *k*-mers-based approach on the quality of the assemblies, we computed the most commonly used metagenome assembly quality metrics, i.e., N50 length, total assembly length, number of contigs, and the maximum contig length [31]. The average N50 length (20.70±9.1 Kbp), assembly size (78466.50±29750.66 Kbp), the total number of contigs (10875.74±5411.13), and the maximum contig length (391.78 ±113.46 Kbp), for the gut metagenomes, obtained by the reduced *k*-mers set was comparable (Wilcoxon rank sum test, *P* >0.05) with negligible differences with the other two sets (Fig.1(b)-1(e), Supplementary Table III). In contrast, we observed that the extended set performed comparatively better than the other two sets by generating a larger assembly size (83797.53±32221.23 Kbp), greater N50 (21.56±9.45 Kbp), and greater maximum contig size (397.36±107.39 Kbp). However, the assembly produced by the extended set was less contiguous and computationally expensive as it took approximately 3X more processing time compared to the reduced *k*-mers (Fig.1(a)-1(e), Supplementary Table III).

For skin metagenomes, the trend remained the same i.e., the reduced *k*-mers set proved equally efficient as it produced comparable quality assembly with average total length (4495.19±7980.70 Kbp), greater N50 (9.9±13.84Kbp), number of contigs (869.33±1789.18), and maximum contig length (99.61±141.77 Kbp) to that of default and the extended sets. However, the extended set produced relatively better results with greater assembly size (5200.0±9413.6 Kbp), and greater maximum contig (112.36±159.01 Kbp) but remained significantly more expensive on time (Wilcoxon rank sum test *P* value <0.0001). Results are shown in Supplementary Fig. 1(b)-1(e). Skin samples were assembled using the reduced *k*-mers set in processing times that were half (3±2.43 min) as compared to the default (7.4±5.78 min) and quarter as compared to the extended (11.75±8.95 min) *k*-mers sets, respectively (Supplementary Fig.1(a)).

The assessment of contig quality using DeepMAsED indicated the presence of a similar fraction of misassembled contigs in the gut metagenome assemblies generated using the three *k*-mers sets. It is worth mentioning here that, from the initial pool of contigs, the lowest fraction (∼19.8%) of contigs was removed from assemblies generated by the reduced *k*-mers set due to low coverage. In contrast, ∼23.6% and 24.6% of contigs were removed due to low coverage from default and extended *k*-mers sets, respectively, by DeepMAsED before predicting misassemblies. Next, we used different thresholds of misassembly scores (MAS) to evaluate the rates of misassemblies among the three *k*-mers sets. First, using a flexible MAS threshold of ³0.2, we observed that none of the contigs was classified as misassembled in the three groups. By using a moderately stringent threshold of MAS<0.1, we observed misassembled contig fractions of 5.1%, 5.3%, and 5.1% in default, reduced, and extended *k*-mers sets, respectively. Applying an extremely stringent threshold (MAS <0.05), the fractions of misassembled contigs increased to 35.3%, 37.6%, and 35.1% in the three *k*-mers sets, respectively (Table I).

**TABLE I:** Comparison of the fraction of misassembled contigs from selective gut metagenomes.

| Sample | *k*-mers | Total<br>Contigs | Assessed by<br>DeepMAseD | %Mis-<br>assembled<br>(*MAS>0.1) | %Mis-<br>assembled<br>(*MAS>0.05) |
| --- | --- | --- | --- | --- | --- |
| SRR1952509 | Default | 12092 | 9113 | 5.9 | 32.3 |
|  | Reduced | 11159 | 8866 | 6.8 | 37.1 |
|  | Extended | 12076 | 8975 | 6.1 | 31.6 |
| SRR1952521 | Default | 5172 | 3769 | 5.0 | 34.5 |
|  | Reduced | 4885 | 3769 | 5.2 | 38.5 |
|  | Extended | 5178 | 3724 | 4.4 | 34.4 |
| SRR1952582 | Default | 10281 | 8300 | 5.2 | 37.0 |
|  | Reduced | 10455 | 8818 | 5.4 | 37.6 |
|  | Extended | 10286 | 8185 | 5.2 | 36.5 |
| SRR2175726 | Default | 10164 | 7732 | 4.1 | 30.8 |
|  | Reduced | 9836 | 7715 | 4.7 | 33.7 |
|  | Extended | 10186 | 7707 | 4.2 | 30.8 |
| SRR2175764 | Default | 6584 | 5391 | 3.2 | 27.2 |
|  | Reduced | 6036 | 5150 | 3.3 | 29.3 |
|  | Extended | 6734 | 5438 | 3.2 | 27.1 |
| SRR2175766 | Default | 11556 | 8620 | 4.9 | 33.8 |
|  | Reduced | 11569 | 9067 | 4.6 | 33.9 |
|  | Extended | 11661 | 8597 | 4.7 | 32.8 |
| **SRR2175768** | Default | 21379 | 16664 | 5.0 | 36.2 |
|  | Reduced | 20996 | 17223 | 5.0 | 38.2 |
|  | Extended | 21459 | 16414 | 5.1 | 36.2 |
| **SRR2241305** | Default | 10359 | 8188 | 5.2 | 34.8 |
|  | Reduced | 9739 | 7896 | 6.0 | 38.2 |
|  | Extended | 10482 | 8173 | 5.2 | 34.5 |
| **SRR2241409** | Default | 12675 | 9801 | 6.4 | 45.4 |
|  | Reduced | 12690 | 10508 | 6.0 | 46.1 |
|  | Extended | 12794 | 9809 | 6.5 | 45.9 |

### Improved Quality MAGs Recovered from Assemblies with Reduced k-mers

Next, we sought to perform the recovery of MAGs from the respective metagenome assemblies of the three sets of *k*-mers. For this, we employed two of the most used genome binning tools, i.e., MetaBAT2 and MaxBin 2.0. For the gut metagenome assemblies obtained using the reduced *k*-mers set, 1983 and 1739 MAGs were recovered using the two binning tools, respectively. The default set generated 2126 with MetaBAT2 and 1765 MAGs with MaxBin 2.0 (Supplementary Table IV-V). The extended set, however, only marginally performed better than the other two sets by producing 2206 and 1831 MAGs from the two tools, respectively. In contrast, as expected, skin samples yielded fewer MAGs across all *k*-mers sets using the two tools (Supplementary Table IV-V).

The raw count of MAGs is not essentially an appropriate metric of choice for describing the quality or determining the utility of the recovered genomes for downstream analyses. We, therefore, deployed CheckM for estimating the completeness and contamination levels of the MAGs recovered from the recruited metagenomes using both tools (MetaBAT2 and MaxBin2.0) across all *k*-mers sets. Interestingly, the MAGs recovered from the gut metagenomes assembled with the reduced *k*-mers set were comparatively less contaminated and more complete. With MetaBAT2, our reduced *k-*mers set generated MAGs with a mean completeness level of 57.52±37.7% and contamination levels of 6.57±24.48%, in contrast with completeness and contamination levels of 56.49±38.05% and 6.66±24.72%, and 55.86±38.42% and 7.11±24.99%, using the default and extended sets, respectively (Fig.2 (a)-2(c)). Utilizing MaxBin 2.0, reduced *k*-mers again produced comparably more complete (68.31±30.73%) and less contaminated (11.96±15.46%) MAGs as compared to the default (completeness: 69.85±29.92%, contamination: 12.84±16.33%) and the extended set (completeness: 68.9±30.38%, contamination 12.47±15.44%) (Fig.2(d)-2(f)). Similarly, for skin metagenomes, the reduced *k*-mers set yielded MAGs that were more complete (MetaBAT2: 59.28±31.88%, MaxBin 2.0: 75.04±28.21%) and less contaminated (MetaBAT2: 4.10±16.28%, MaxBin 2.0: 11.30±16.24%) when compared to the other sets (Supplementary Fig.2(a)-2(f) and Supplementary Table IV-V).

**Figure 2:**
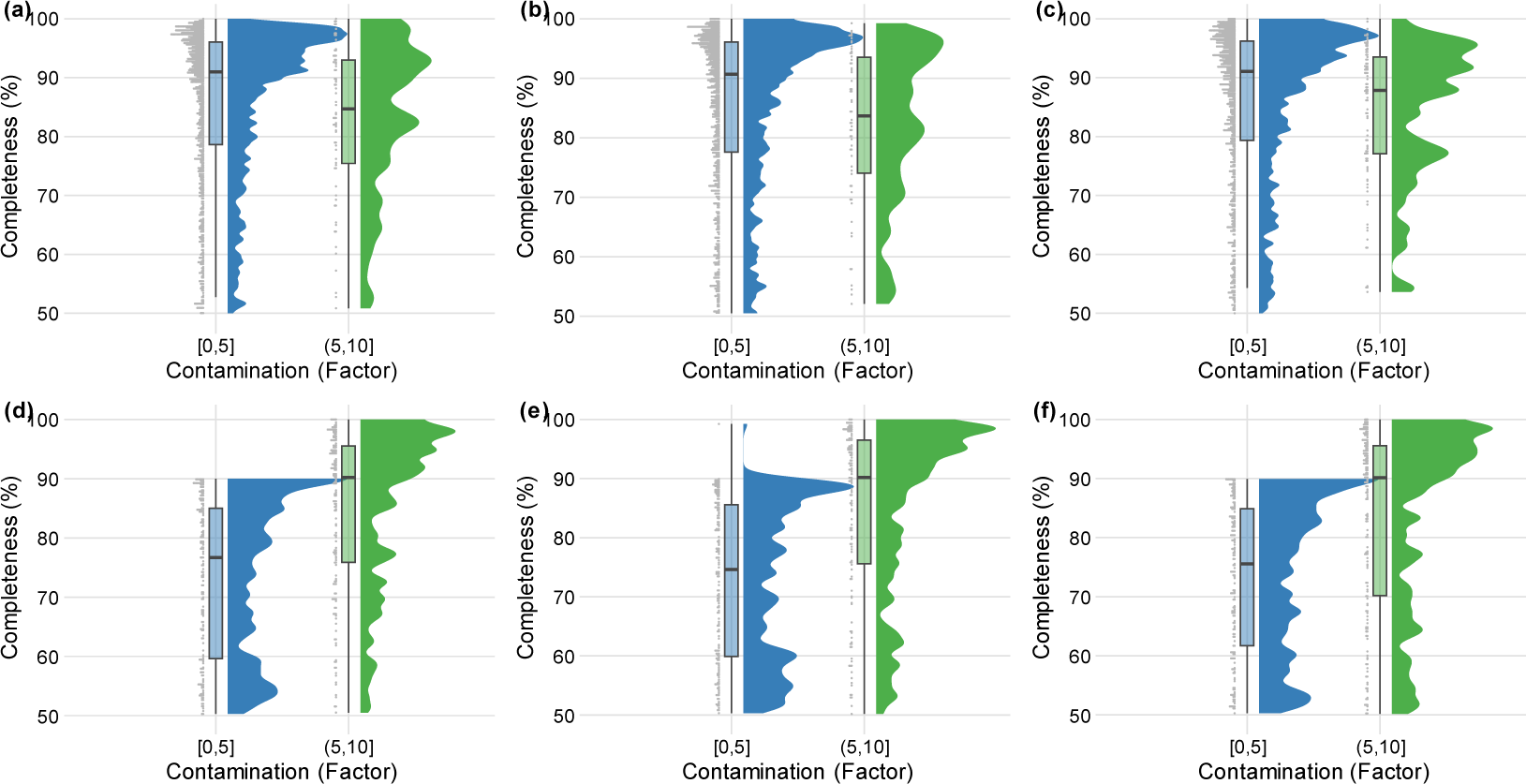
Distribution of MAGs recovered from human gut metagenome samples across all three *k*-mer sets as per their contamination and the completeness levels. (a) Distribution of MAGs with completeness above 50% and contamination range of 5-10% obtained via MetaBAT2 using the Default set. **(b).** MAGs distribution acquired by the reduced *k*-mer set using MetBAT2. **(c).** Distribution of MAGs obtained by employing the extended *k*-mer set using MetaBAT2. **(d).** Distribution of MAGs obtained by the application of Default *k-*mer set with MaxBin 2.0 **(e).** Distribution of MAGs recovered with the reduced *k*-mer set using MaxBin 2.0 **(f).** Distribution of MAGs acquired with the extended *k*-mer set by employing MaxBin 2.0.

We further categorized the recovered MAGs into HQ, MQ, and LQ groups based on the completeness and contamination cutoffs defined in the MIMAG criteria (see Methods for more details). This categorization indicated that relatively higher proportions of HQ and MQ MAGs were recovered from the assemblies generated by the reduced *k*-mers set. For gut metagenomes, among the MAGs recovered using MetaBAT2, 25.7%, and 27.0% were categorized as HQ and MQ, respectively. In contrast, the default set yielded 25.4% HQ and 25.3% MQ MAGs, while the extended set produced 23.8% HQ and 24.7% MQs, respectively. Furthermore, the reduced *k*-mers approach decreased the proportion of LQ MAGs by 1%. Similar outcomes were obtained for the MAGs recovered with MaxBin 2.0 (Table II). Finally, the reduced *k*-mers set re-exhibited promising results for the skin metagenomes by generating a 3% higher proportion of both HQ and MQ MAGs using MetaBAT2 and MaxBin 2.0 (Supplementary Table IV-V, respectively).

**TABLE II:** Proportion of HQ, MQ, and LQ MAGs recovered from gut samples using MetaBAT2 and MaxBin 2.0.

| **MetaBAT2** |  |  |  |
| --- | --- | --- | --- |
| *k*-mers set | Default | Reduced | Extended |
| MAG Count | 2126 | 1983 | 2206 |
| HQ% | 25.4 | 25.7 | 23.9 |
| MQ% | 25.3 | 27.0 | 24.8 |
| LQ% | 48.4 | 47.3 | 51.3 |
| **MaxBin 2.0** |  |  |  |
| *k*-mers set | Default | Reduced | Extended |
| MAG Count | 1765 | 1739 | 1831 |
| HQ% | 19.7 | 20.2 | 20.2 |
| MQ% | 20.9 | 19.0 | 20.3 |
| LQ% | 59.3 | 60.7 | 59.5 |

### Computational Time Efficiency with The Reduced k-mers Set

After conducting a thorough quality assessment, we analyzed the total processing time to recover genomes from the metagenomic samples. This total time was the aggregated real-time duration of each sub-process, including assembly, indexing, read mapping and sorting, estimation of contigs depths, genome binning, and quality assessment of MAGs. Notably, all the analyses were conducted on Ubuntu 22.10 (GNU/Linux 5.19.0-46-generic x86_64) with 48 threads and 150 GB RAM. A statistical comparison of the total time across different *k*-mers sets proclaimed the reduced *k*-mers approach as optimal.

On average, using MetaBAT2, the processing time for gut metagenomes was 47.0±10.9 min, significantly reduced from the default and extended set, which required 61.4±18.4 min, and 102.5±28.2 min, respectively (Fig.3(a)). For MaxBin 2.0, the gut samples consumed ∼47.9±10.2 min with the reduced *k-*mers set as compared to default (61.6±17.8 min) and the extended (103.02±28.0 min) sets (Fig.3(b). The statistical differences, tested using the Wilcoxon rank sum test, declared these differences in the processing times as highly significant with *P* <0.0001 with both MetaBAT2 and MaxBin 2.0 (data not shown). In contrast, skin metagenomes understandably required significantly less processing time i.e. ∼8.31±6.62 min with MetaBAT2 and 8.45±6.36 min for MaxBin 2.0 as compared to the default (12.7±9.3 min) and extended (17.27±12.3 min) sets (Supplementary Fig.3(a), 3(b)) and Supplementary Table VI).

**Figure 3:**
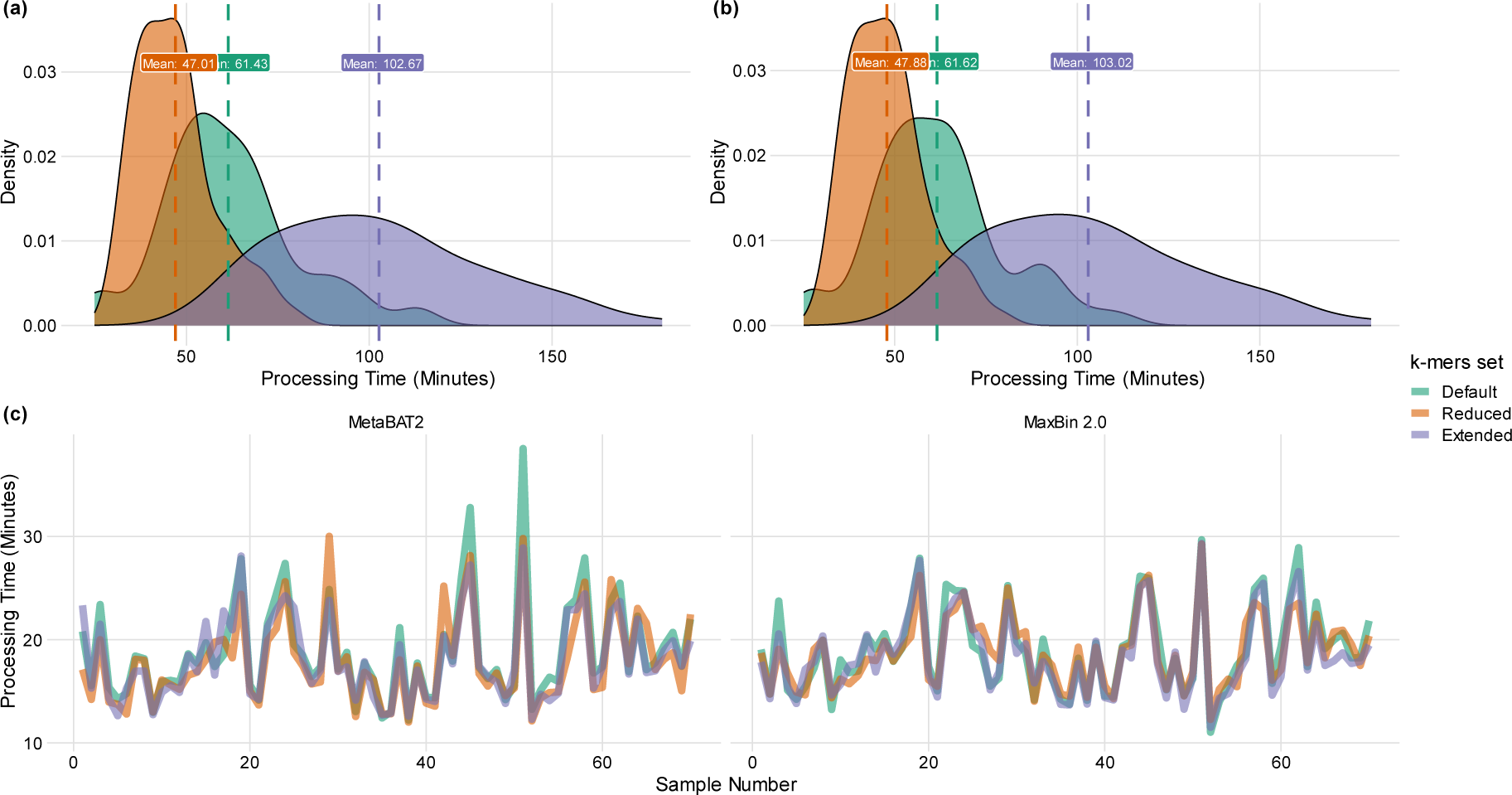
Comparison of the total time taken by gut samples. (**a)** Total processing time taken by the samples from assembly to MAGs QC using MetaBAT2 across all three *k-*mer sets **(b)** Sum of the times taken by the sub-processes i.e., from assembly to MAGs QC using MaxBin2.0 across default, reduced and extended *k*-mer sets. **(c).** Comparative analysis of the total time consumed by sub-processes except assembly using all three *k*-mer set with MetaBAT2 and MaxBin 2.0 respectively.

Furthermore, if the *de novo* assembly time is excluded from the total processing time, both MetaBAT2 and MaxBin 2.0 required similar processing times for gut and skin metagenomes for the other analysis steps (Fig. 3(c) and Supplementary Fig. 3(c)).

### The Reduced k-mers Set Outperformed the Results Obtained by Pasolli et al. 2019

For further validation of the utility of the reduced *k*-mers set, we selected a subset of gut metagenomes (Supplementary Table II) described previously by Vincent C et al. [29]. These metagenomes were assembled with MEGAHIT using default parameters, whereas MetaBAT2 was used for recovering MAGs in a large-scale study on the human metagenomes conducted by Pasolli et al. [1]. *De novo* metagenome assembly and recovery of MAGs for these metagenomes using the reduced *k*-mers set yielded significantly better results than Pasolli et al.

The qualitative comparison was performed employing the metagenome assembly properties, including the total length, number of generated contigs, size of the largest contig, N50, and percentage of the GC content. In line with our prior findings, these results again affirmed our reduced *k*-mers set as an optimal choice for better results. The assembly lengths yielded in our study (39695.49±21341.80 Kbp) were similar to what was achieved in Pasolli et al. (42162±22653.09 Kbp) as shown in Fig.4(a). Our approach assembled these metagenomes with a significantly higher N50 length (Wilcoxon rank sum test *P* <0.01). The mean N50 in this study (19.25±23.01 Kbp) nearly doubled when compared to the N50 lengths (9.87±7.46 Kbp) for these metagenomes obtained by Pasolli et al. (Fig.4(d)). Comparison between the lengths of the largest contig produced in both studies indicated that our approach, with the maximum contig size of 298±141.29 Kbp, surpassed (Wilcoxon rank sum test, *P* <0.001) largest contig length obtained by Pasolli et al. (178.83±79.55 Kbp) (Fig.4(c)). Lastly, using our approach, the metagenomes were assembled with marginally better contiguity (6785.52±4288.56 Kbp) than Pasolli et al. (8541.18±5416.16 Kbp), however, these differences were not statistically significant (Fig.4(b), Supplementary Table VII).

**Figure 4:**
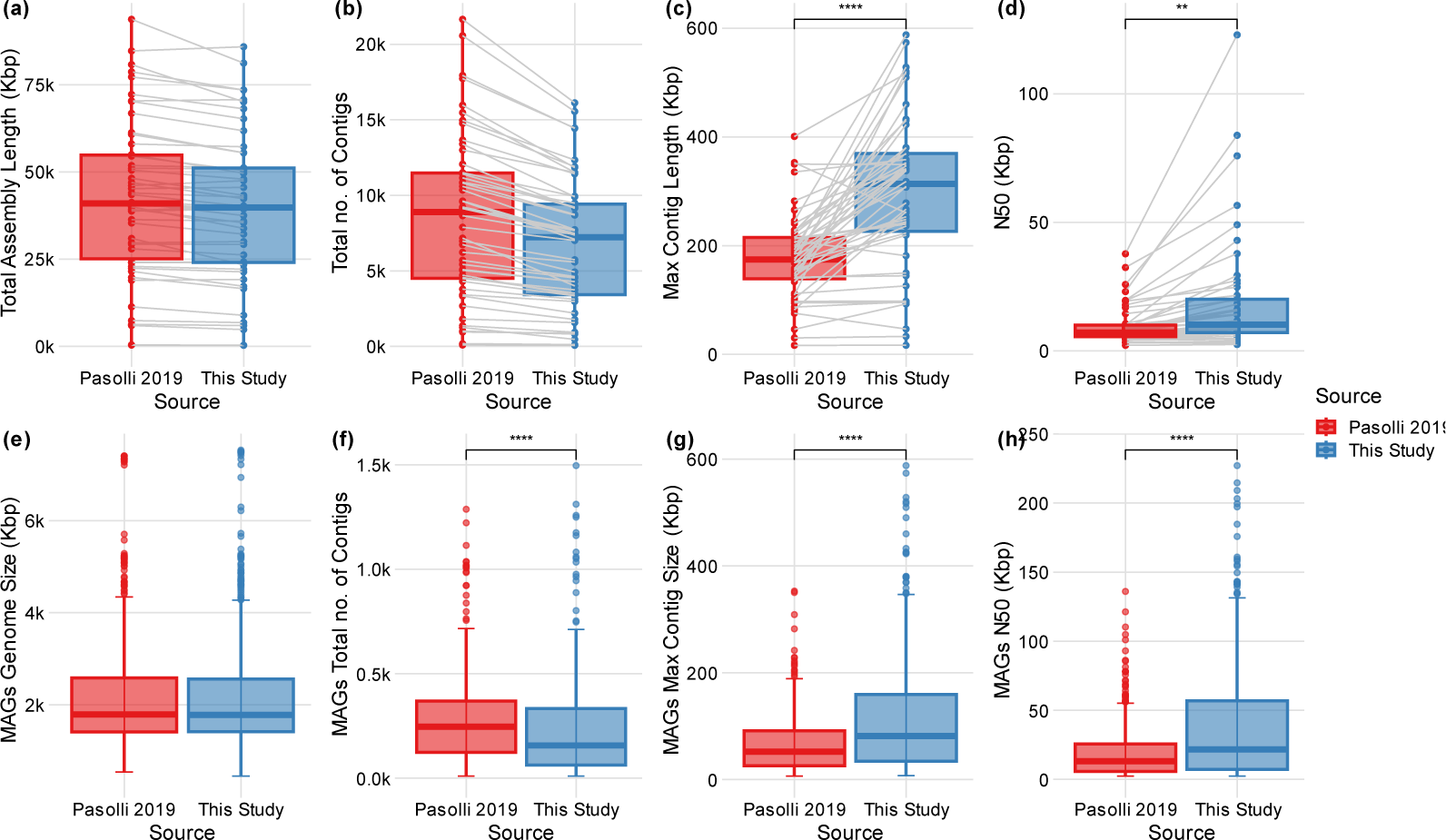
Comparison between the assembly results achieved by Pasolli et al. using MegaHit’s default configurations and this study’s reduced *k*-mer approach. **(a)** Comparison between the total assembly lengths generated by both studies. **(b)** Comparison between the total number of contigs attained by the two approaches **(c)** Comparison between the maximum contig size produced by Pasolli et al. and this study. **(d)** Comparison between the N50 as achieved by Pasolli et al. and this study. **(e)** Comparison between the total genome sizes of MAGs as acquired by Pasolli et al. and this study. (**f)** Total number of contigs contained by MAGs as yielded by both studies. **(g)** Comparison between the maximum contig length of MAGs obtained by Pasolli et al. and this study. **(h)** N50 length comparison between the MAGs yielded by the two studies. **Note:** In the figure, asterisks represent the p-value as determined by the Wilcoxon rank sum test (** = <0.01, *** = <0.001 and ****= p-value <0.0001).

Finally, we sought to determine the differences between the quality of MAGs recovered from these metagenomes belonging to the two groups. Our approach resulted in a relatively higher number of MAGs than Pasolli et al. (450 vs. 445). Further analyzing the categorization of these MAGs, indicated that our method yielded a higher proportion of HQ MAGs (48.89%), accompanied by a relatively lower occurrence of MQ MAGs (51.11%) in contrast with 46.74% HQ and 53.26% MQ MAGs yielded by using default *k*-mers set by Pasolli et al. (Supplementary Table VIII).

Furthermore, Pasolli et al. obtained a larger genome size (2837.84±1629 Kbp) for HQ MAGs, than genome sizes for HQ MAGs obtained using the reduced *k*-mers set (2812.575±1623.52 Kbp). However, their genomes were contained in a higher number of contigs (224.11±163.03 vs 155.79±169.74). By comparing other metrics, we observed further improvements in MAGs recovered using our approach. For instance, HQ MAGs had a drastically improved N50 (53.41±45.56 Kbp) that was ∼2X better than HQ MAGs recovered by Pasolli et al. Similarly, the maximum contig size (160.84±118.21 Kbp) was also significantly better (90.05±51.88 Kbp). For the MQ MAGs, the comparison showed almost similar results except that the average genome size of MQ MAGs, generated by the reduced *k*-mers appeared comparatively greater (1793±982.62 Kbp) than Pasolli et al. (1785.33±929.21 Kbp) while the trends of improved contiguity (295.49±262.4 vs Pasolli’s 336.03±232.84), double N50 length (25.33±36.13 Kbp vs 13.98±19.74 Kbp) and larger maximum contig size (72.56±78.82 Kbp vs 47.45±48.5 Kbp) remained consistent (Fig.4(e)-4(h), Supplementary Table VIII).

## DISCUSSION

Choosing the right parameters for the available bioinformatics tools is one of the most crucial tasks that can directly affect the quality as well as the accuracy of the results. Fine-tuning these parameters can enable the users to obtain reliable results in reduced time as well as within the limited computational resources. In DBG-based metagenome assemblers, the most crucial parameter is *k,* which determines the size of *k*-mers to fragment the reads. If the *k* value is smaller, it can tangle the graph and produce more fragmented reads, while the larger values of *k* can yield erroneous assemblies [18], [32]. Determining the optimal *k* requires an extensive amount of memory and processing time, which becomes practically impossible in certain scenarios [33]. Hence, the decision to choose *k*-mers presents a critical trade-off between meta-assembly quality and computational cost. Based on these observations, we hypothesized that a set of right *k*-mers could resolve the issue by producing improved assembly quality in a limited time. Therefore, in this study, we tested two sets, a reduced set with fewer *k* values but a larger step size (i.e., 28) and an extended set with more *k* values but a smaller step size (i.e., 6) to explore the effects and come up with an optimal *k*-mers settings, in addition to the default set of *k*-mers.

Here, we used MEGAHIT, which is an ultrafast and efficient metagenome assembler as compared especially with metaSPAdes [34] and is commonly employed in genome-centric metagenome analyses with default parameters [1], [35], which we believe is not an optimal option, especially in computational resource-limited settings. Hence, we empirically designed an optimal configuration that can achieve maximum computational efficiency with the production of assembly and genomes of the highest quality.

Consequently, our results have demonstrated that the reduced *k*-mers set can perform exceptionally well for human microbiome metagenome analyses performed on DBG-based assemblers. This approach can improve the results by producing contiguous assemblies while cutting the required computational time (taken by default *k*-mers) to half. The insignificant variations observed in the metagenome assembly metrics, as obtained by the reduced *k*-mers approach, align with our goal of maintaining the assembly integrity while optimizing the computational cost. The negligible differences in assembly length, N50, number of contigs, and the maximum contig length could be viewed as a trade-off in the pursuit of improved speed and resource utilization without significantly compromising the overall assembly quality.

Assembly quality is often assessed by comparing the re-sequenced genome with its reference genome. In the case of metagenomes, contigs are typically assembled *de novo*, thus the assessment of the quality of the metagenome assembly becomes a very challenging task. MetaQUAST [36], uses closely related reference genomes for the detection of misassemblies (translocation, inversion, and relocations) from the metagenomic contigs. However, this approach can be satisfactorily applied to a limited number of previously sequenced species. Therefore, we used DeepMAsED, which deploys deep learning to estimate misassemblies and is an ideal choice for performing a reference-independent quality assessment of the metagenomic contigs. It must be noted here that the accuracy of DeepMAsED can decrease if the contig is longer than 10 Kbp. We purposely selected the metagenomes that produced longer contig lengths using the reduced *k*-mers set and expected a significantly greater fraction of misassemblies in the contigs assembled using the reduced *k*-mers set. However, despite this, and after using stringent MAS thresholds, we observed only negligibly higher misassemblies (0.2% and 2.3%, respectively) in the contigs obtained using the reduced *k*-mers and the other two sets. The 2.3% higher misassemblies can be attributed to a highly stringent MAS threshold (MAS<0.05) used as a cutoff or to the inherent limitation of DeepMAsED in handling contigs larger than 10 Kbp. Nonetheless, the misassembled contigs could be easily identified from the outputs produced by DeepMAsED and excluded from downstream analyses.

Exploiting the reduced *k*-mers set for the genome binning produced a lesser number of MAGs as compared to the default settings; however, this reduction can be interpreted as a balance between quantity and quality as the MAGs recovered by the reduced *k*-mers exhibited higher completeness and lower contamination level, especially with MetaBAT2. Additionally, it is worth mentioning that comparatively less contaminated MAGs were recovered using MetaBAT2 whereas MaxBin

2.0 produced MAGs with better completeness. Moreover, MetaBAT2 resulted in a relatively greater number of MAGs than MaxBin 2.0 for all three *k*-mers sets. For further assessing the quality of MAGs, we employed the MIMAG criteria by considering the contamination and completeness thresholds while excluding the evaluation of proportions of 5S, 16S, 23S rRNA, and tRNAs required for classifying a MAG as HQ. This decision was influenced by the nature of our work which did not require gene annotation. Here, the reduced *k*-mers resulted in the recovery of relatively higher proportions of HQ and MQ MAGs accompanied by lower proportions of LQ MAGs, affirming that the reduced *k*-mers can serve as an optimal solution. This supports that, regardless of the choice of tools used for MAG recovery, the reduced *k-*mers can yield higher-quality MAGs.

Lastly, validation of results by applying the reduced *k*-mers set on the dataset from Pasolli et al. further attested this approach as significantly optimal. By this comparison, it is evident that the reduced *k*-mers set has the potential to produce desirably contiguous assemblies with significantly reduced computational time. This was supported by a lower number of contigs, larger maximum contig sizes, and N50 values in the relevant dataset. Moreover, more complete and less contaminated MAGs, recovered from assemblies generated using the reduced *k*-mers, further assert that using default *k*-mers may not be the most ideal solution in large-scale genome-centric metagenomic analyses.

The ability of reduced *k-*mers set to generate contiguous assemblies more rapidly (∼2X faster) will provide the platform to revolutionize genome-centric studies. This development will facilitate the rapid identification of compositional and functional potentials of microbiomes, characterization of the properties of microbial communities, and extraction of near-complete genomes from metagenomes. Considering the results presented above, we highly recommend the adoption of reduced *k*-mers settings as an optimal solution for a time-efficient assembly of the human metagenome samples as well as recovery of MAGs from the assemblage. Our comprehensive analyses validate the improved performance of this approach, both in terms of efficient metagenome assembly and recovery of MAGs. We believe that its implementation will significantly expedite the research endeavors of the scientific community while optimizing the computational cost.

While this study has made significant contributions by identifying the reduced *k-*mers approach tailored for the complex human microbiome samples, we must acknowledge certain limitations. Firstly, our analysis was conducted empirically and lacked a systematic approach for *k*-mers selection due to the limited availability of the relevant literature on this concept. Next, it is important to note that at some points of the analysis, the extended *k*-mers set produced superior in terms of assembly contiguity, genome recovery rates, and the proportions of HQ MAGs. However, this improvement came at the cost of a 3X increase in the processing time. This disparity underscores the vast potential for further improvements in *k-*mers subset optimization and indicates the feasibility of further optimizing the sets of *k-mers* that can achieve an even finer balance between producing high-quality results and conserving limited computational resources.

## CONCLUSION

This study is focused on human microbiome metagenomes, similar investigations must also be extended to other metagenomic datasets along with the exploration of other pertinent parameters. The systematic exploration of the parameters employed in the existing metagenomic analysis tools holds the potential to significantly contribute to addressing the computational constraints and improving the quality of results.

## ACKNOWLEDGMENT

We are highly thankful to the Supercomputing Lab at the School of Interdisciplinary Engineering and Sciences, NUST, for providing the necessary computational resources for this work.

