## Supplementary material for "Efficient *De Novo* Assembly and Recovery of Microbial Genomes from Complex Metagenomes Using a Reduced Set of *k*-mers": Figure 1-3

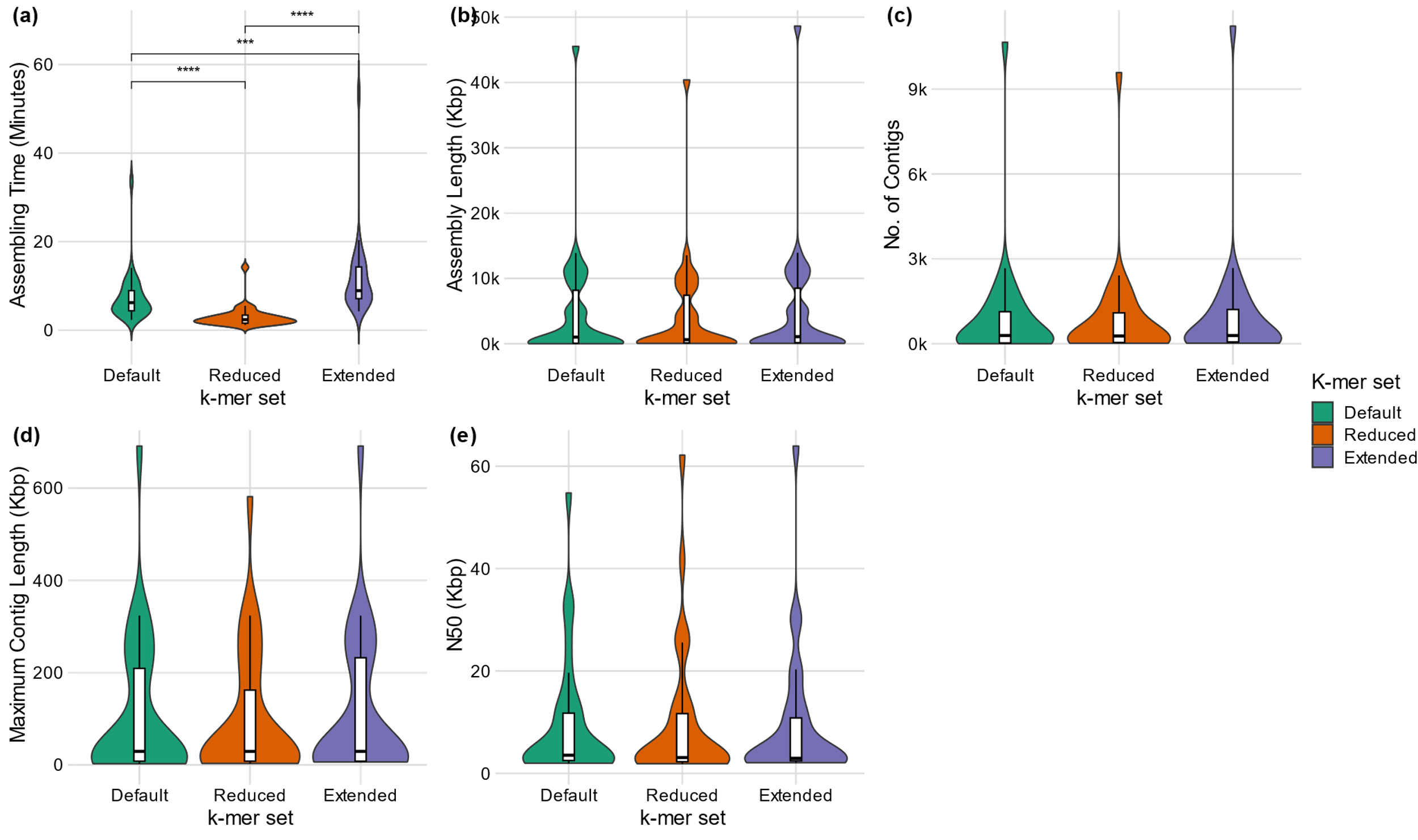

**Fig.1.** **Comparative analysis of the assembly metrics of human skin metagenome samples between default, reduced and extended *k*-mer sets.** (**a)** Comparison in the processing time of the assembly performed using different *k*-mer sets. **(b)** Pairwise comparison between the number of contigs generated by the three *k*-mer sets. **(c)** Comparison between the N50 lengths of the assembly attained by different *k*-mer configurations. **(d)** Comparison between the total assembly length achieved by the different *k*-mers sets. **(e)** Comparison between the maximum contig length acquired by the three *k*-mer sets. **Note:** In the figure, asterisks represent the p-value as determined by the Wilcoxon rank sum test (** = <0.01, *** = <0.001 and ****= p-value <0.0001).

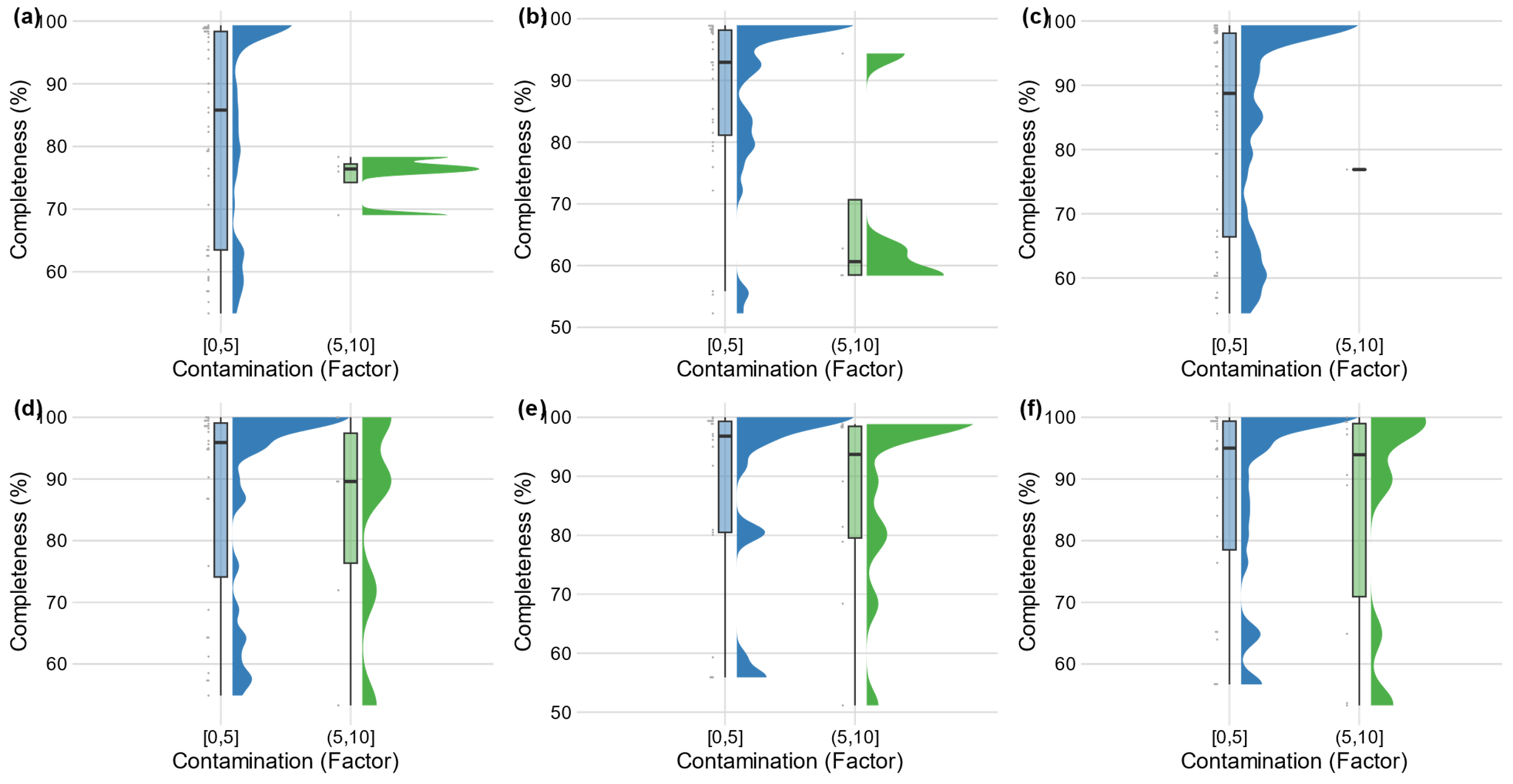

**Fig. 2. Distribution of MAGs recovered from human skin metagenome samples across all three *k*-mer sets as per their contamination and the completeness levels (a)** Distribution of MAGs with obtained via MetaBAT2 using the Default set. **(b).** MAGs distribution acquired by the reduced *k*-mer set using MetBAT2. **(c).** Distribution of MAGs obtained by employing the extended *k*-mer set using MetaBAT2. **(d).** Distribution of MAGs obtained by the application of Default *k-*mer set with MaxBin 2.0 **(e).** Distribution of MAGs recovered with the reduced *k*-mer set using MaxBin 2.0 **(f).** Distribution of MAGs acquired with the extended *k­*-mer set by employing MaxBin 2.0.

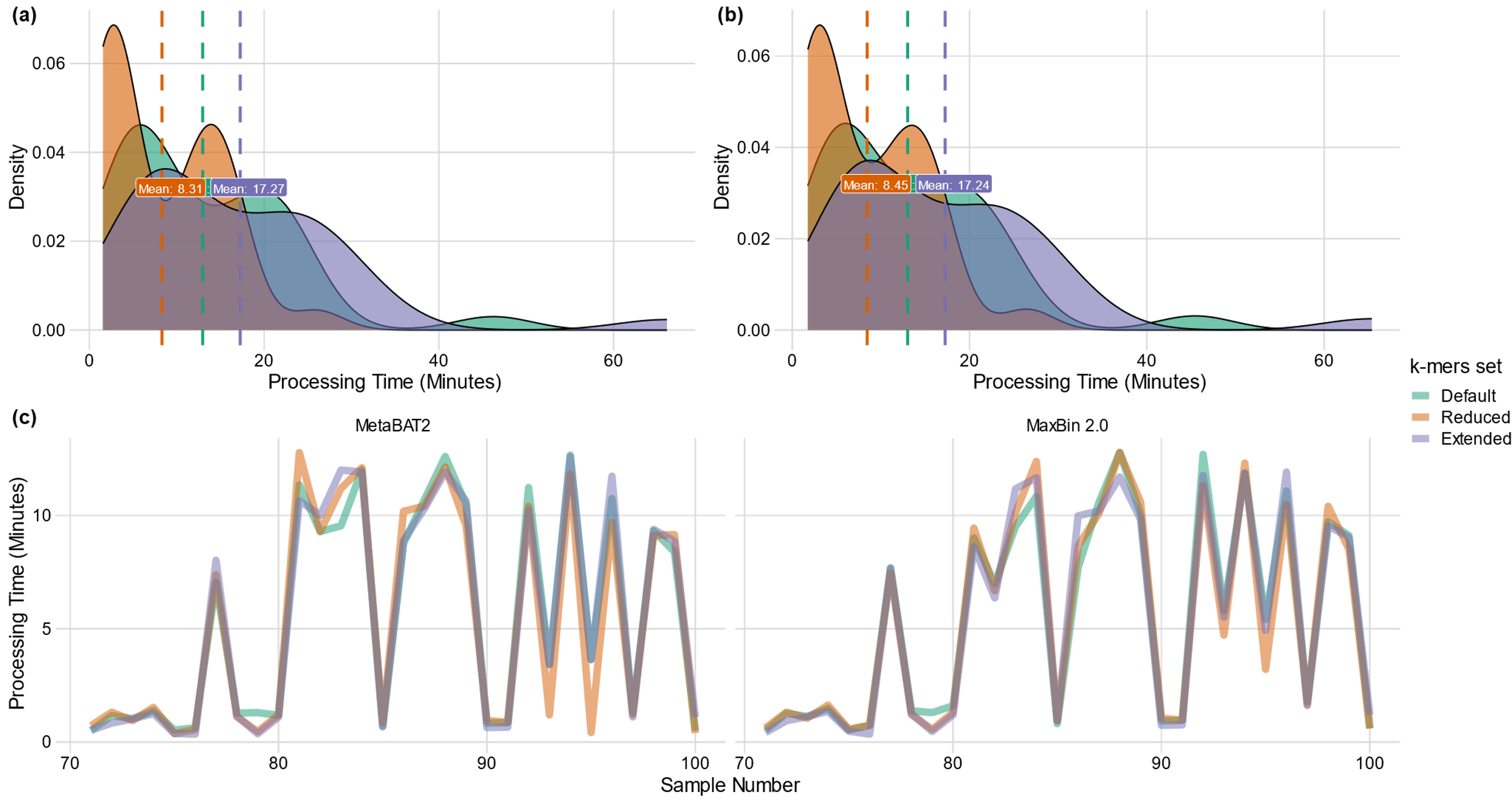

**Fig.3: Comparison of the total time taken by gut samples.** (**a)** Total processing time taken by the samples from assembly to MAGs QC using MetaBAT2 across all three *k-*mer sets **(b)** Sum of the times taken by the sub-processes i.e., from assembly to MAGs QC using MaxBin2.0 across default, reduced and extended *k*-mer sets. **(c).** Comparative analysis of the total time consumed by sub-processes except assembly using all three *k*­-mer set with MetaBAT2 and MaxBin 2.0 respectively.
